# Managing beach access and vehicle impacts following reconfiguration of the landscape by a natural hazard event

**DOI:** 10.1101/2022.02.15.480598

**Authors:** Shane Orchard, Hallie S. Fischman, David R. Schiel

**Affiliations:** Marine Ecology Research Group, University of Canterbury, Private Bag 4800, Christchurch 8140, New Zealand; School of Earth and Environment, University of Canterbury, Private Bag 4800, Christchurch 8140, New Zealand; Department of Environmental Engineering Sciences, Engineering School of Sustainable Infrastructure and Environment, University of Florida, Gainesville, Florida 32611, USA

**Keywords:** nature-based recreation, recreational access, shoreline management, impact assessment, disaster recovery, protected areas

## Abstract

After New Zealand’s 7.8 Mw Kaikōura earthquake in late 2016 an unexpected anthropogenic effect involved increased motorised vehicle access to beaches. We show how these effects were generated by landscape reconfiguration associated with coastal uplift and widening of high-tide beaches, and present analyses of the distribution of natural environment values in relation to vehicle movements and impacts. Access changes led to extensive vehicle tracking in remote areas that had previously been protected by natural barriers. New dunes formed seaward of old dunes and have statutory protection as threatened ecosystems, yet are affected by vehicle traffic. Nesting grounds of nationally vulnerable banded dotterel (*Charadrius bicinctus bicinctus*) co-occur with vehicle tracking. An artificial nest experiment showed that vehicle strikes pose risks to nesting success, with 91% and 83% of nests destroyed in high and moderate-traffic areas, respectively, despite an increase in suitable habitat. Despite gains for recreational vehicle users there are serious trade-offs with environmental values subject to legal protection and associated responsibilities for management authorities. In theory, a combination of low-impact vehicle access and environmental protection could generate win-win outcomes from the landscape changes, but is difficult to achieve in practice. Detailed information on sensitive areas would be required to inform designated vehicle routes as a potential solution, and such sensitivities are widespread. Alternatively, vehicle access areas that accommodate longstanding activities such as boat launching could be formally established using identified boundaries to control impacts further afield. Difficulties for the enforcement of regulatory measures in remote areas also suggest a need for motivational strategies that incentivise low-impact behaviours. We discuss options for user groups to voluntarily reduce their impacts, the importance of interactions at the recreation-conservation nexus, and need for timely impact assessments across the social-ecological spectrum after physical environment changes -- all highly transferable principles for other natural hazard and disaster recovery settings worldwide.

## 1. Introduction

Off-road vehicles (ORV) present many possibilities for accessing remote areas where there are no formed roads. However, the potentially adverse effects of these activities create challenges for environmental managers who must identify the existence, extent and severity of associated impacts, and design appropriate responses. Moreover, vehicular access can also facilitate other valued activities such as camping, hunting and fishing, leading to the need to consider the relative merits of vehicular access in various forms and its relationship with established objectives. This study investigates changes to vehicular beach access generated by the reconfiguration of a natural landscape. These changes were caused by the 7.8 M_w_ Kaikōura earthquake that struck the east coast of New Zealand in November 2016, affecting over 130 km of coastline spanning two local government jurisdictions. The area included a c. 40 km section of the Marlborough region characterised by extensive sandy and mixed sand-gravel beaches that are the focus of this study. This relatively remote area is known for its wild and scenic values and has few road access points (Figure 1).

**Figure 1.**
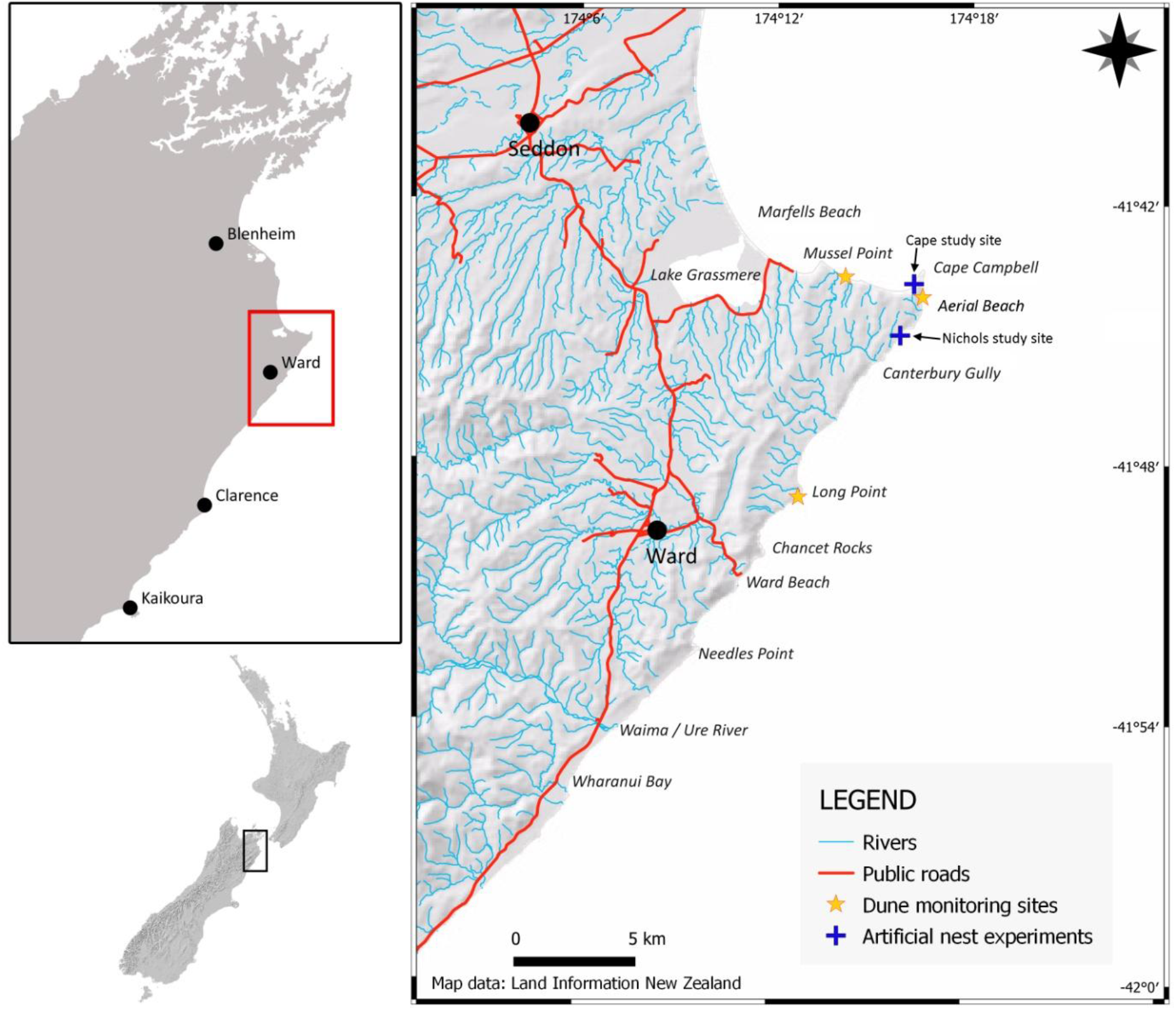
Overview of the study area on the east coast of the South Island of New Zealand.

The earthquake caused a series of complex ruptures and fault movements associated with a highly variable pattern of vertical displacement (Hamling et al. 2017; Holden et al. 2017; Xu et al. 2018). At the coastline, this displacement was mostly uplift by as much as 6 m (Clark et al. 2017; Orchard et al. 2021), leading to the widening of beaches. Soon after the earthquake, vehicle movements along the coast increased as an anthropogenic side-effect of topographical changes that improved access opportunities. These changes represent initially un-noticed and un-managed human responses with an apparent gain for recreational vehicle users, but with potentially undesirable aspects that remained unexplored. Moreover, there are transferable potential lessons for other large-scale landscape changes in disaster recovery and resource management contexts elsewhere.

In many parts of the world, ORV access is reportedly increasing and in many cases their cumulative effects have not been adequately addressed or managed. For example, Priskin (2003) reported that the number of ORV access points had increased by 115% between 1965 and 1998 on the Central Coast of Western Australia. Drivers may also drive along the edge of existing formed tracks, widening the area of disturbance (Davenport & Davenport 2006). Studies from Queensland have reported more concentrated vehicle traffic on the upper beach, with the lower beach being used to varying degrees when exposed on lower tides (Schlacher & Thompson 2007).

The severity of ORV impacts depends on factors such as the frequency, timing and type of traffic, composition and vulnerability of biological communities, and the precise location of vehicle tracking in relation to sensitive areas. Several studies on ORVs have reported detrimental impacts on the structure and function of sandy beach ecosystems. For example, beaches with ORV traffic may have dunes set further back from the high tide line than beaches closed to ORVs (Houser et al. 2013), indicating impacts on the formation of foredunes in areas they would otherwise occupy. Direct impacts of ORV traffic include sand compaction, erosion and vegetation loss leading to dune destabilisation (Davenport & Davenport 2006; Groom et al. 2007; Hosier & Eaton 1980; Houser et al. 2013). Because dunes act as a barrier against storm surge and sea-level rise, damage to them can also reduce protection for housing and other infrastructure to landward (Orchard & Schiel 2021). The potential for damage to plant communities is an important consideration for ORV impact assessments because of the sand-trapping properties of dune plants (Martinez et al. 2016). The vegetation-mediated impacts of ORVs are potentially more important than direct erosion effects of vehicle tracking since much larger volumes of sand may be released from dune systems by wind erosion in the absence of plant cover. In New Zealand, where invasive marram grass (*Ammophila arenaria*) has displaced native sand-binders, ORV impacts may also contribute to the demise of native plant communities and their associated fauna, thereby creating an additional conservation issue (Stephenson 1999).

Shorebirds are among the most studied wildlife in relation to ORV impacts because of the potential vulnerability of key life stages such as nesting. Crushing by vehicles is particularly problematic for species that lay camouflaged eggs in cryptic nests such as ‘scrapes’ on beaches (Stephenson 1999; Weston et al. 2012). Disturbance by vehicles may also cause birds to reduce their feeding time or increase time away from the nest (Defeo et al. 2009). Studies in Queensland found that only 34% of ORV drivers slowed down or changed course when they encountered a shorebird, and that partial beach closures were effective in reducing the rate of egg crushing (Weston et al. 2012; Weston et al. 2014). Additionally, numerous studies have shown adverse ORV impacts on beach invertebrates, including crustaceans and shellfish that are vulnerable to crushing (Davies et al. 2016; Lucrezi & Schlacher 2010; Moss & McPhee 2006; Schlacher et al. 2007; Schlacher et al. 2008a).

Methods to control ORV impacts include the use of non-regulatory signage in sensitive areas (Supplementary Material Figure S1). Additionally, statutory measures may be imposed through legal instruments such as bylaws. These have been implemented for the control of vehicle impacts on several beaches in New Zealand to date, although are often hotly contested by stakeholders. In this case study, post-disaster responses for coastal management have included the development of a proposed beach vehicle bylaw (Marlborough District Council 2021), and related aspects will require integration within longer-term planning arrangements. To support these initiatives our research programme included a suite of social-ecological investigations across a spatially extensive coastal area. Our objectives included the quantification of beach profile changes, vehicle tracking patterns, dune system responses and distribution of nesting grounds for banded dotterel (*Charadrius bicinctus bicinctus*) as an indicator species currently listed as ‘nationally vulnerable’ in the New Zealand Threat Classification System (Robertson et al. 2017).

## 2. Methods

### 2.1 Shoreline characteristics and change

The study area is a contiguous 36 km section of the Marlborough coastline stretching from the Waima / Ure River in the south to Marfells Beach in the north (Figure 1). This area marks the northern extent of coastal uplift associated with the Kaikōura earthquake. The degree of uplift in the study area was in the 1-3 m range and mainly associated with movements on the Needles Fault (Clark et al. 2017; Hamling et al. 2017; Holden et al. 2017). The coastal environment in this area differs considerably from the predominantly rocky Kaikōura coast further south and includes extensive sandy beaches and dune systems interspersed with headlands. Coastal reef substrates include relatively soft sedimentary rock platforms than have experienced high rates of erosion (Schiel et al. 2019). The combination of unique geology and a generally dry climate are associated with high faunal and floristic diversity and the coast is an important area for migratory bird species (de Lange et al. 2013; Jones & Hutzler 2002; Marlborough District Council 2021).

Pre-quake shorelines for the study area were digitised for the spring high tide line and dune system toe on the high tide beach using visible markers such as vegetation limits, strand lines and drying heights visible in high resolution aerial photography acquired 2015. This imagery was comparable to immediate post-quake data acquired two years afterwards, with the pixel size of both datasets being 0.2 m (Table 1). Assessment of the pre-quake landscape was also assisted by comparison with a 2002 aerial imagery dataset that happened to be captured at high tide, creating a useful reference for inundation levels at many points along the coast (Table 1). Additionally, the whole area was surveyed on foot in 2018 and 2019 and GPS coordinates captured for a wide range of landmarks and reference points including pre-quake vegetation zonation patterns and the position of new recruitment zones. Contour intervals at 0.1 m were extracted from the post-quake LiDAR and the 0.6 m contour was adopted as a representative position for the estimated high tide line based on the abovementioned visual clues. This vertical elevation is equivalent to a mean high water spring tidal height of ~0.4m NZVD and an additional 0.2 m (vertical) swash zone (Land Information New Zealand 2021).

**Table 1.** Aerial imagery and LiDAR datasets

(a) Aerial imagery
| Dataset | Acquisition date | Supplier | Commissioning agency | Imagery specifications |  |
| --- | --- | --- | --- | --- | --- |
|  |  |  |  | Resolution | Format |
| Marlborough 1m Rural Aerial Photos (2002) | Various dates in 2002 | Terralink International | Marlborough District Council | 100 cm pixel | Black and White GeoTIFF |
| Marlborough 0.2m Rural Aerial Photos (2016) | 7 January 2016 - 7 March 2016 | AAM NZ Ltd | Marlborough District Council | 20 cm pixel | 3-band (RGB) GeoTIFF |
| Post-quake Aerial Imagery<sup>†</sup> | 3 December 2016 - 6 January 2017 | AAM NZ Ltd | Land Information New Zealand | 20 cm pixel | 3-band (RGB) GeoTIFF |

(b) LiDAR data
| Dataset | Acquisition date | Supplier | Commissioning agency | Accuracy specification (m) |  |
| --- | --- | --- | --- | --- | --- |
|  |  |  |  | vertical | horizontal |
| Post-quake Airborne LiDAR<sup>†</sup> | 3 December 2016 - 6 January 2017 | AAM NZ Ltd | Land Information New Zealand | ±0.10 | ±0.50 |

<sup>†</sup> captured concurrently

A set of 720 cross-shore (perpendicular to the shoreline) transects were generated at 50 m intervals along a smoothed baseline prepared from the pre-quake high tide line using the *ambur* package in R (Jackson et al. 2012). Intersection analyses in a GIS environment were used to compute cross-shore beach width and shoreline changes in relation to the timing of the earthquake (Figure 2). To examine spatial co-occurrence patterns, a complementary set of belt transects, each 50 m wide at the baseline, was generated in a northward direction from the first transect origin at Waima River in the south. Substrate types for the belt transects were classified using field data according to four classes (reef, boulder, mixed sand-gravel, sand) that represent the predominant substrate on the high tide beach. Collectively, these belt transects covered the entire coast and were of a sufficient cross-shore dimension to intersect with all other field survey and remote-sensed data as described in the sections below.

**Figure 2.**
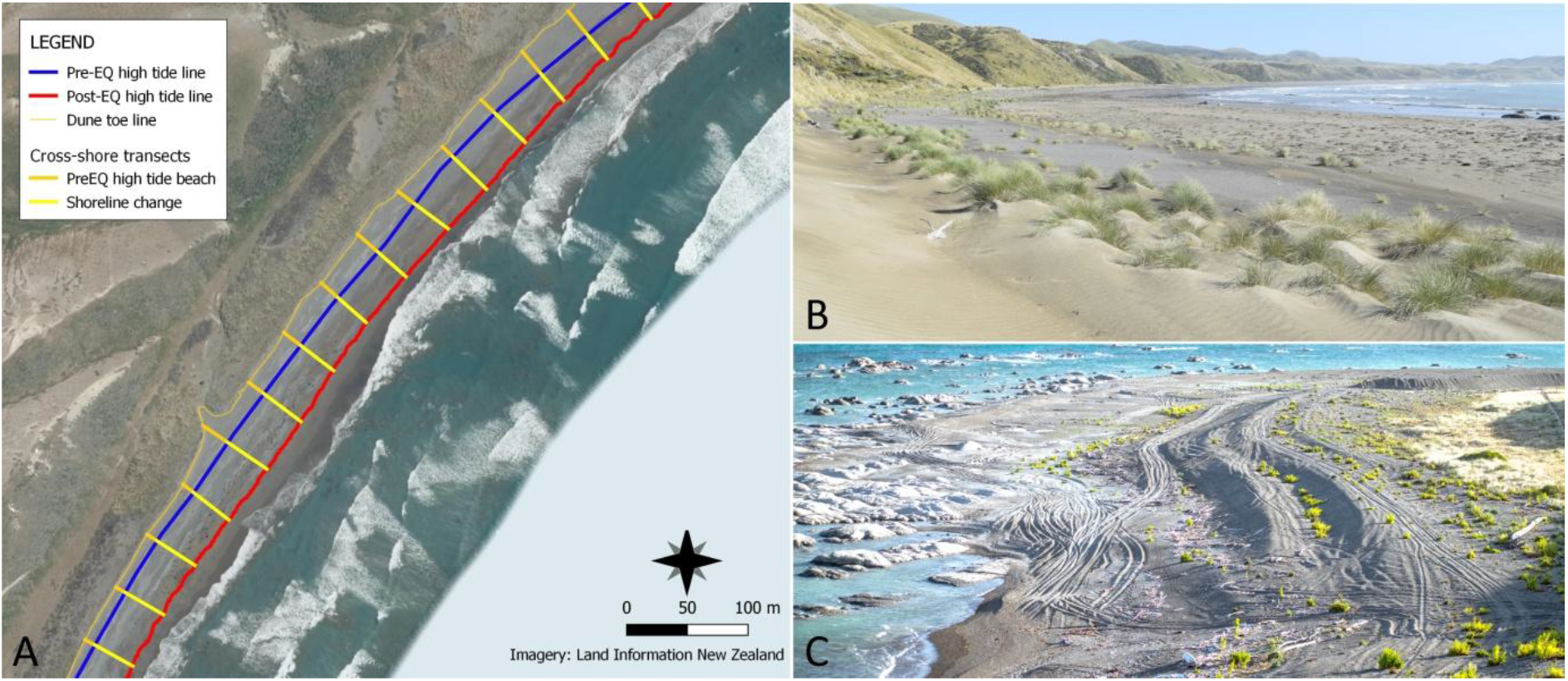
(A) Example of the shoreline change sampling set-up showing cross-shore transects at 50 m spacing and two example calculations overlaid on post-quake aerial imagery. The area in (A) is shown in (B) located north of Long Point where the beach has widened considerably and new dunes have formed. (C) Post-quake vehicle tracking on the high tide beaches between Cape Campbell and Canterbury Gully.

### 2.2 Off-road vehicle tracking

Off-road vehicle (ORV) tracking measurements were made at periodic intervals throughout the study area and included whole-coast surveys completed in the summers of 2018 and 2019. The width of visible vehicle tracks was measured in the cross-shore direction to the nearest metre. Monitoring points included at the location of changes in vehicle tracking patterns and at the position of all shorebird nesting territories (described below). This measurement reflects the distance between the tyre tracks of individual vehicles or, in the case of overlapping tracks, the dimension between the outer tyre marks, summed across the shore profile. Only tracking above the high tide line was considered, to avoid biases introduced by observations of tracking below the high tide line which are influenced by survey timing in relation to the tide. Additional information collected included evidence of preferred routes, such as where tracks were seen to converge or fan out in response to barriers and topographic changes, and the location of access points and turnaround areas.

### 2.3 Shorebird nesting sites

#### Census surveys

Surveys of banded dotterel nesting territories were completed for the entire coastline in November 2018 and 2019. Coordinates were recorded using handheld GPS receivers for the estimated midpoint of nesting territories based on bird behaviour. Visual clues used to identify territories included characteristic ward-like behaviour, which includes the ‘escorting’ of intruders to beyond the territorial range, in addition to distraction behaviour such as broken wing displays that are indicators of a nest or chicks nearby. In areas supporting several nesting territories the boundaries were discriminated using two observers following the movements of adjacent breeding pairs. Additional searches were made for nests at two locations (Ward Beach and Canterbury Gully) to assess the cross-shore location of nests on the high tide beach with a focus on establishing whether newly created habitat associated with beach widening was being utilised for nesting sites. Nesting territory mid-points were overlaid on the belt transects to assess relationships with substrate types and vehicle tracking density.

#### Artificial nest experiment

To provide a more direct test of the risk of vehicle strikes, an artificial nest experiment was conducted at two sites after the breeding season. These experiments used quail eggs which are very similar to banded dotterel eggs in size and colouration and were deployed in shallow scrapes to provide a close representation of banded dotterel nests (Supplementary Material, Figure S2). The Cape Campbell site represented a location with a high traffic density of c.70% tracking on a relatively narrow high tide beach 20-25 m wide. The Nicholls site represented a location with a moderate traffic density of c.40% tracking on a high tide beach 60 m wide. At each site, six representative territories were established, each occupying 50 m in the long-shore direction. To examine the potential consequence of nesting locations in old vehicle tracks (which is relative common for banded dotterel), potential nest sites within each territory were stratified according to their previous tracking status (tracked / untracked). Four nests were placed at random locations within each territory using a 1 m grid and applying a minimum distance to any vehicle track of 2 m to define the untracked class. Additionally, a buffer of 5 m minimum was maintained from the upper and lower limits of the high tide beach to reduce confounding edge effects. Both study sites were mixed sand-gravel beaches and the area around each nest was smoothed to facilitate the detection of predator tracks.

All nests were monitored weekly for nest failures which were classified according to five classes of threats (vehicle strike, trampling by horses, avian predation, small mammal predation, and unknown / other predation) based on visual clues. Vehicle strikes were readily identifiable from recent tracking and the observation of crushed nests, as was the single example of trampling by horses. Avian predation was evidenced by the observation of peck marks with the egg shell being typically crushed inwards (Supplementary Material, Figure S2). Small mammal predation was recorded at a few nests where it was associated with partially eaten eggs. Other predation was the class assigned to the disappearance of eggs, often accompanied by mammalian tracks being observed in the vicinity of the nest. These losses are likely attributable to larger and more vigorous predators such as mustelids or feral cats. As the data followed a non-normal distribution, Kruskal Wallis tests were used to assess the effects of territory and previous tracking status on nest losses at each study site. To incorporate the effect of site within the analysis and improve statistical power using the combined dataset we fitted Generalized Liner Mixed Models with binomial error distributions to both total nest losses and nest losses by vehicle strikes. In both GLMMs, we treated time (week) as a random effect and previous tracking status, site, and their interaction as fixed effects. Analyses were produced in R version 4.0.3 (R Core Team 2021), using the *tidyverse* package (Wickham et al. 2019), and glmer function in the *lme4* package for the GLMMs (Bates et al. 2008).

### 2.4 Dune system responses

Field transects were established at three sandy beaches located east of Mussel Point, south of Cape Campbell (Aerial Beach) and south of Long Point (Figure 1). Each of these beaches supports the native sand-binder spinifex (*Spinifex sericeus*) and the post-quake response of spinifex dunes is of considerable interest for beach conservation objectives. At each monitoring site, two cross-shore transects were established from an origin point in the back dune swale. Each transect spanned the pre-quake foredune and newly developing dune system on the uplifted beach. The Mussel Point and Long Point sites featured spinifex remnants in the old dune system that were expected to colonise the new accommodation space created by uplift through the vegetative growth of runners from established plants. At the Aerial Beach site there were no spinifex remnants in the old dune system due to extensive invasion by marram grass along this section of coast, but several relatively large spinifex patches had become established on the high tide beach (likely reflecting recruitment soon after the earthquake). Measurements taken in the summers of 2018 and 2019 included the canopy cover and maximum height of vegetation within contiguous 2 x 2 m plots along the transect line that together provided a complete description of the vegetation composition within a 2 m belt across the dune profile. Canopy cover estimation followed the method of Hurst & Allen (2007) which considers the area within the perimeter of the canopy of each plant as seen in plan-view for each species of interest. Bare ground cover was recorded where it occurred and vehicle tracking measured as detailed above. Ground elevation was measured at 1 m intervals along each transect using a combination of laser level and real-time kinematic (RTK) GPS surveys with a Trimble R8 GNSS receiver. Geodetic benchmarks were included within all RTK-GPS surveys and referenced to the New Zealand Vertical Datum (NZVD) (Land Information New Zealand 2016), to achieve an estimated vertical accuracy of 3.5mm ± 0.4 ppm RMS.

## 3. Results

### 3.1 Shoreline change

Seaward movement of the pre-quake high tide line was observed on all transects although the magnitude of the changes was highly variable, as expected, due to variable uplift (Figure 3A). Shoreline change ranged from 1.2 m - 205 m with a mean of 30.6 ± 0.7 m (SE). Prior to the earthquake 90% of the coastline featured high tide beaches that were <50 m wide. This percentage was halved to 45% as a result of beach widening induced by coastal uplift. The mean cross-shore width of high tide beaches increased from 25.6 ± 0.9 m to 56.2 ± 1.2 m. Across all transects, the minimum width increased from zero to 7 m, and maximum width increased from 180 m to 247 m (Figure 3B).

**Figure 3.**
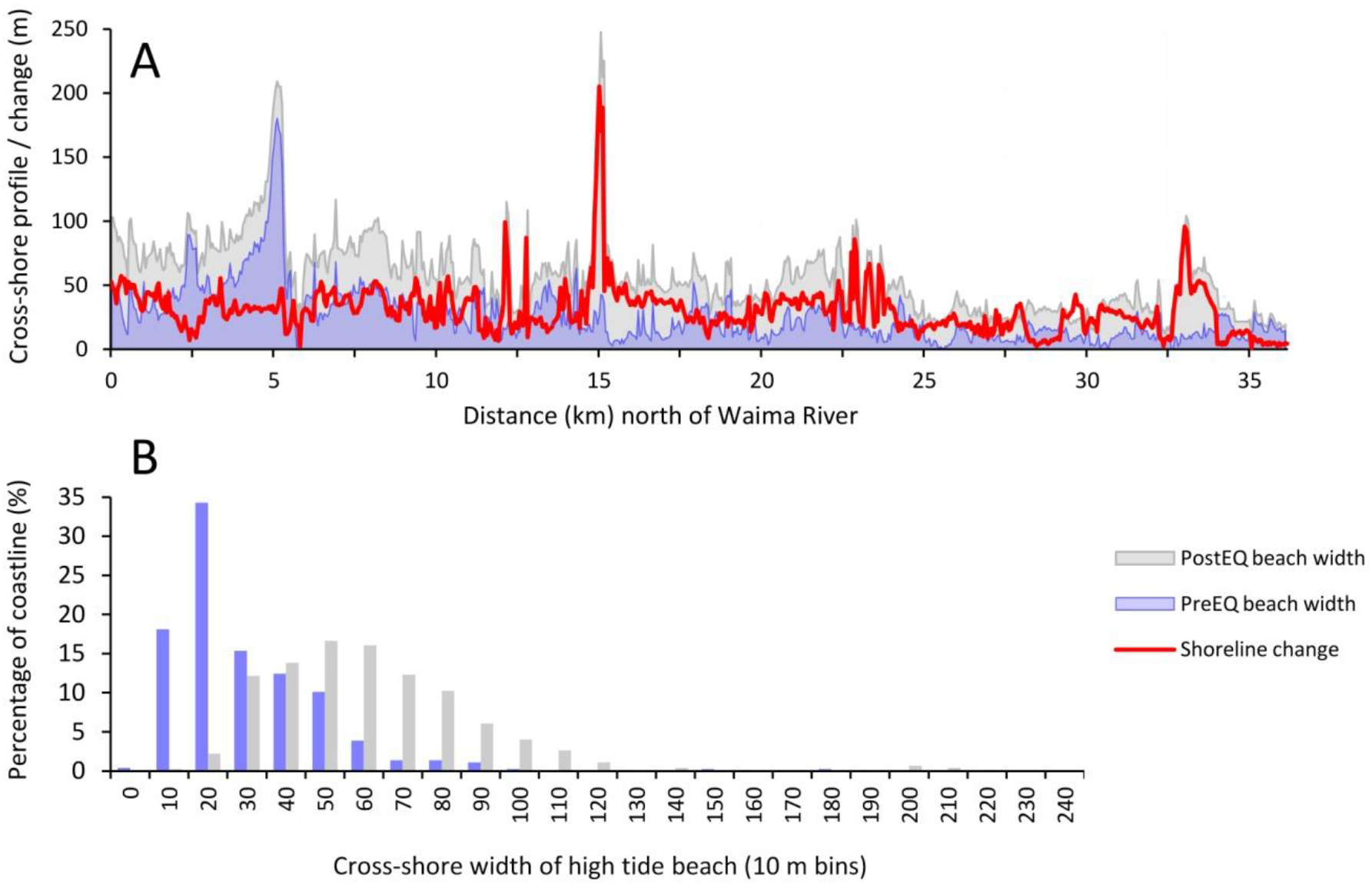
(A) Cross-shore width of the high tide beach before and after the 2016 Kaikōura earthquake along 36 km of coastline from the Waima River in the south to Marfells Beach in the north. (B) Histograms of the high tide beach width calculated for 10 m beach-width increments. The X-axis labels represent the upper value of each 10 m bin. An additional ‘zero width’ bin is also included.

Changes to high tide beaches were considerable. Prior to the earthquake, some sections of the coast had no high tide beach (n=21 transects), while 50 transects (2.5 km of coastline) were <5 m wide, and 157 transects (7.8 km of coastline) were <10 m wide. These narrow sections of coast included rocky headlands at Mussel Point, Cape Campbell, Chancet Rocks and Needles Point (Figure 1). Following the earthquake, there were no transects with high tide beaches <5 m wide and only two <10 m wide, both located at Mussel Point. These results indicate that pre-quake locations with topographical barriers to ORV traffic at high tide had been widened sufficiently to facilitate vehicle access, given that at least 10 m of high tide beach had become available nearly everywhere. These changes are attributable almost solely to uplift of the coastline since the LiDAR data and imagery used to assess post-quake conditions was acquired within a few days of the earthquake.

### 3.2 Vehicle tracking

Vehicle tracking patterns have changed considerably along the coast, although to varying degrees. These are most pronounced at sections of the coastline characterised by steep rocky substrates on the cross-shore profile and steep hill country above that previously created a barrier to most forms of ORV traffic. Changes in the north of the study area (near Marfells Beach) had the greatest influence (Figure 4A). They involved only short sections of rocky coast at Mussel Point and headlands further east that experienced only modest uplift (c. 1m), but it was sufficient to expose a high tide ledges that facilitated all-tide access to Cape Campbell under most combinations of tide and swell. Accompanying changes in ORV usage have included relatively large volumes of traffic (e.g., daily ORVs >30 at Cape Campbell), becoming commonplace in previously remote areas that were seldom visited before.

**Figure 4.**
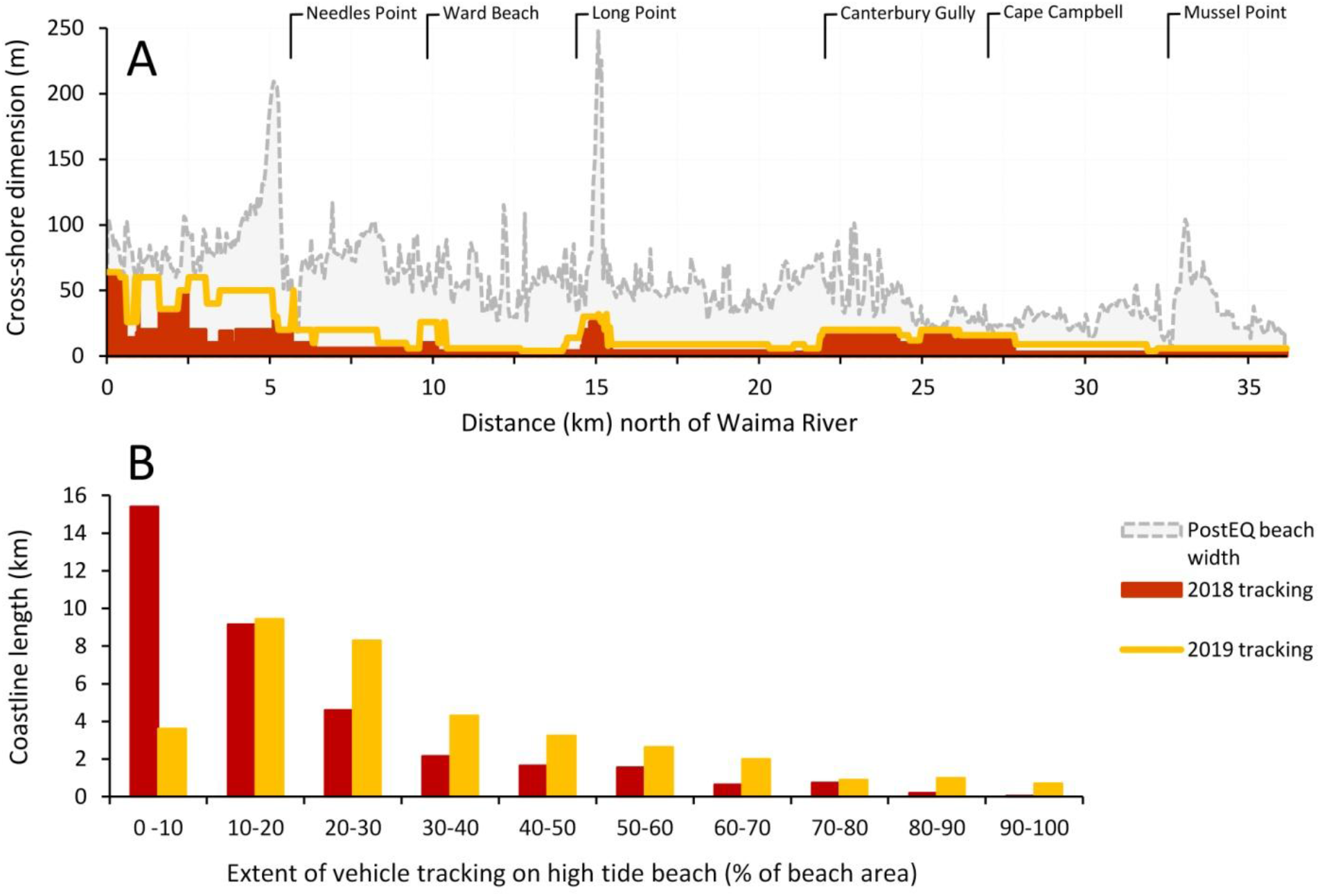
Vehicle tracking patterns across 36 km of coastline between Waima River and Marfells Beach measured over two consecutive summers following the November 2016 Kaikōura earthquake. (A) Cross-shore high tide beach width and extent of vehicle tracking. (B) Histogram of vehicle tracking extent showing the percentage of beach and length of coastline affected.

The vehicle tracking pattern in 2018 showed extensive tracking in the south of the study area between Waima River and Needles Point, but this had also been the case before the earthquake due to the river mouth being a popular vehicle access point (Figure 4A). To the north of Needles Point, tracking was reduced to 6-10 m which is generated mostly by vehicles travelling south from Ward Beach, a popular public access point located 4km further north. Similarly, Chancet Rocks, located 2 km north of Ward Beach, prevented most ORVs from travelling further north from the Ward Beach access point. The stretch of coast immediately north of Chancet Rocks featured the lowest level of vehicle tracking in the study area in both years (equivalent to 1 −2 vehicle tracks). However, many of those vehicle movements originated from Marfells Beach located 25 km to north, in addition to a few private land access points. The cross-shore tracking pattern partly reflects popular route choices to avoid obstacles, such as at Long Point where reef platforms are present lower in the intertidal zone. Between 2018 and 2019 the whole-coast average increased 73% from 10.0 ± 0.4 m to 17.3 ± 0.6 m of tracking on the cross-shore profile, despite the minimum and maximum values staying unchanged at 3 m and 64 m respectively. The associated change in tracking density (taking into account the width of the beach) was a 67% increase from 19.5 ± 0.4% to 32.6 ± 0.8% of the high tide beach area (Figure 4B).

### 3.3 Banded dotterel nesting sites

#### Census survey

A total of 60 nesting territories were identified in the 2018 survey, and 69 territories in 2019 (Figure 5). Although the spatial distribution varied slightly, locations with high nest density were very similar between years. Some of the changes observed were a higher number of nesting pairs south of Cape Campbell on both sides of Canterbury Gully, and lower numbers at Long Point in 2019 in comparison to 2018 (Figure 5). Beaches characterised by mixed sand-gravel substrates supported the majority of nesting territories in both years (73.3% and 76.8% for 2018 and 2019, respectively) (Figure 6A). Territories were also recorded on sandy beaches (21.7% and 17.4% of nesting territories for 2018 and 2019, respectively), in locations such as the bay north of Long Point and at Mussel Point. However, the pattern of substrate fidelity is somewhat blurred by the heterogeneous nature of many beaches that feature patches of various substrate types. On the finer sand beaches, for example, dotterels are still likely to find areas of coarser gravels, which provide more suitable camouflage for the placement of nests.

**Figure 5.**
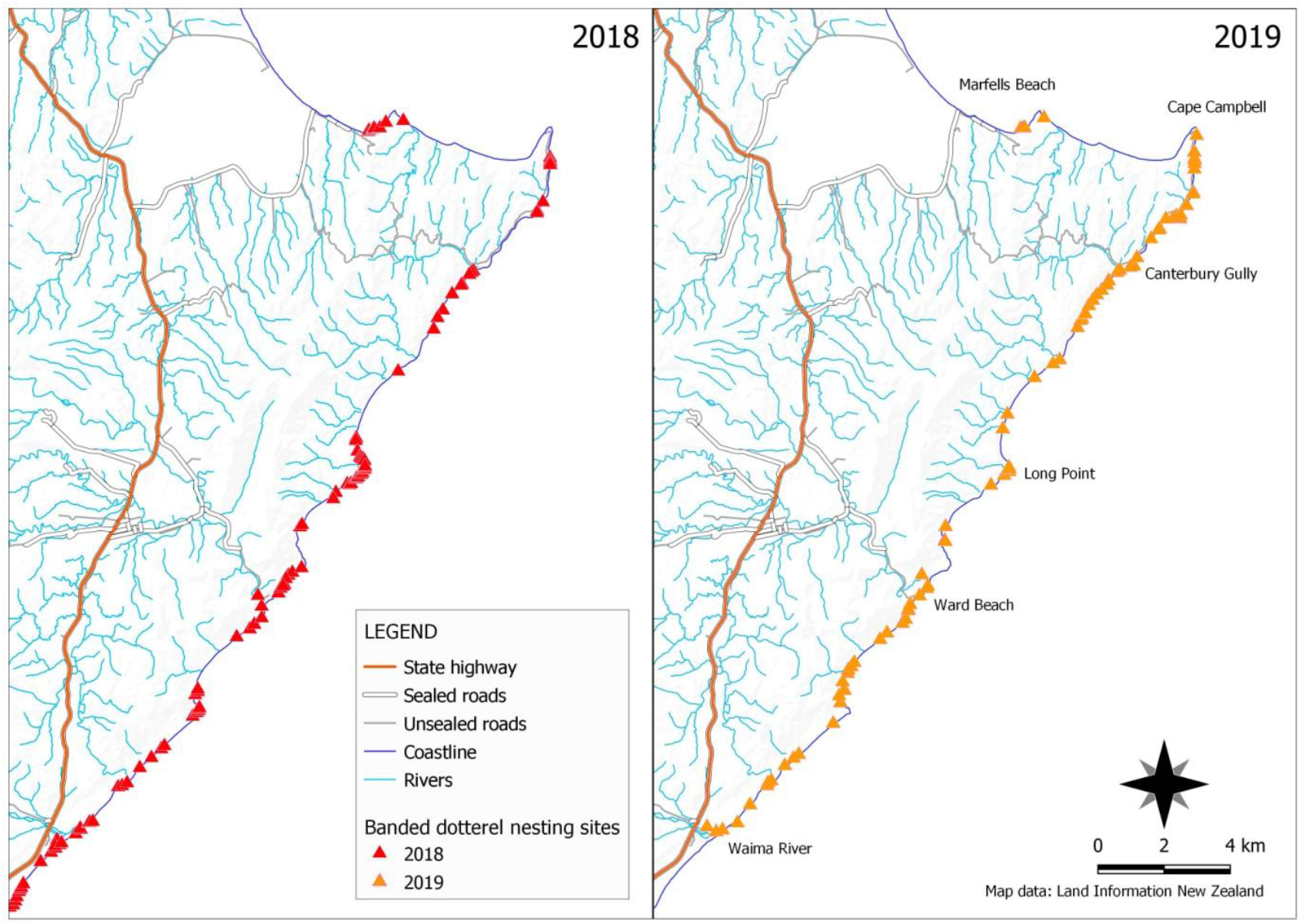
Distribution of banded dotterel (*Charadrius bicinctus bicinctus*) nesting territories across 36 km of coastline between Waima River and Marfells Beach in 2018 and 2019.

**Figure 6.**
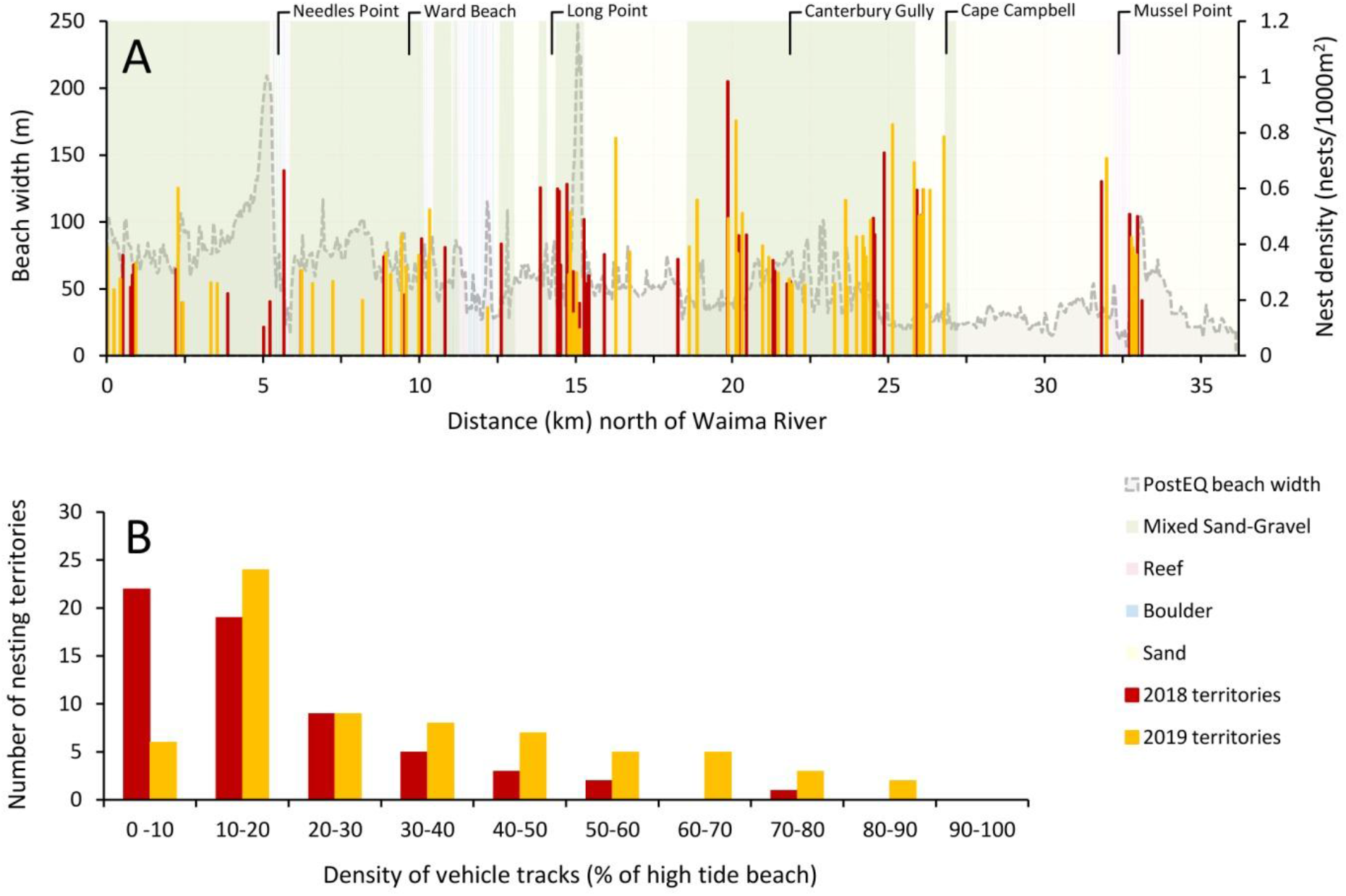
(A) Density of banded dotterel (*Charadrius bicinctus bicinctus*) nesting territories across 36 km of coastline between Waima River and Marfells Beach overlaid on the high tide beach and dominant substrate type. (B) Density of vehicle tracking in nesting territories expressed as percentage of the high tide beach.

Nesting sites may be located anywhere on the high tide beach, including close to the new post-quake high tide line. This indicates that beach widening has increased the area of suitable habitat. All dotterel nesting territories were exposed to some degree of disturbance from motor vehicles, as evidenced by tracking patterns (Figure 6B). The degree of spatial overlap is indicated by the tracking density within dotterel territories which varied by 3–90% in 2018, and 3–99% in 2019. The mean tracking density increased from 19.5 to 32.3% between years within dotterel territories. A histogram of these data shows a decrease in frequency in the 0-10% bin and a relatively consistent increase across other bins, indicative of a gradual increase in the density of tracks in all of the dotterel nesting areas (Figure 6B). It is important to note that these anecdotal tracking patterns record only previous vehicle movements and may include old tracks that persisted from the previous nesting season. Field observations of vehicle movements made on various dates and times suggest that the actual volume of traffic was similar between years.

#### Artificial nest experiment

Over the typical incubation period of four weeks there was a high nest loss rate at both study sites, with 92% of nests failing at the Cape site, and 83% at Nichols (Figure 7). Generalized Linear Mixed Models of total nest losses over time showed that there was a significant difference between the study sites (Z = 2.897, p = 0.004), with the higher tracking density site (Cape) showing a higher nest loss rate (Table 2). This finding is consistent with a correlation between actual vehicle movements during the study period and the previous density of vehicle tracking that was used to characterise and select the study sites. The Cape site also had a higher overall number of vehicle strikes and they occurred earlier in the study period. Vehicle crushing was the single biggest causal factor, accounting for 50% of the nest losses at the Cape site and 40% at Nichols. The GLMMs, however, showed no significant difference between sites for nest losses by vehicle strikes (Z = 1.634, p = 0.102). This partly reflects the smaller difference between sites for the vehicle strike rate versus overall losses, and variance introduced by the previously tracked and untracked nest locations (Figure 7). Previous tracking status was found to be a significant factor for vehicle strikes as the cause of failure (Z = 3.495, p < 0.001), but not for total nest losses (Z = 1.047, p = 0.295). This is consistent with preferential use of existing vehicle tracks by ORVs as expected. Nonetheless, the relatively high nest loss rates at locations that were previously untracked shows that vehicles are continuing to explore new areas. At the Nichols site, 38% of all nests were lost to vehicles on previously untracked parts of the beach, and the situation was worse (46% of nests) with higher traffic density at the Cape study site.

**Figure 7.**
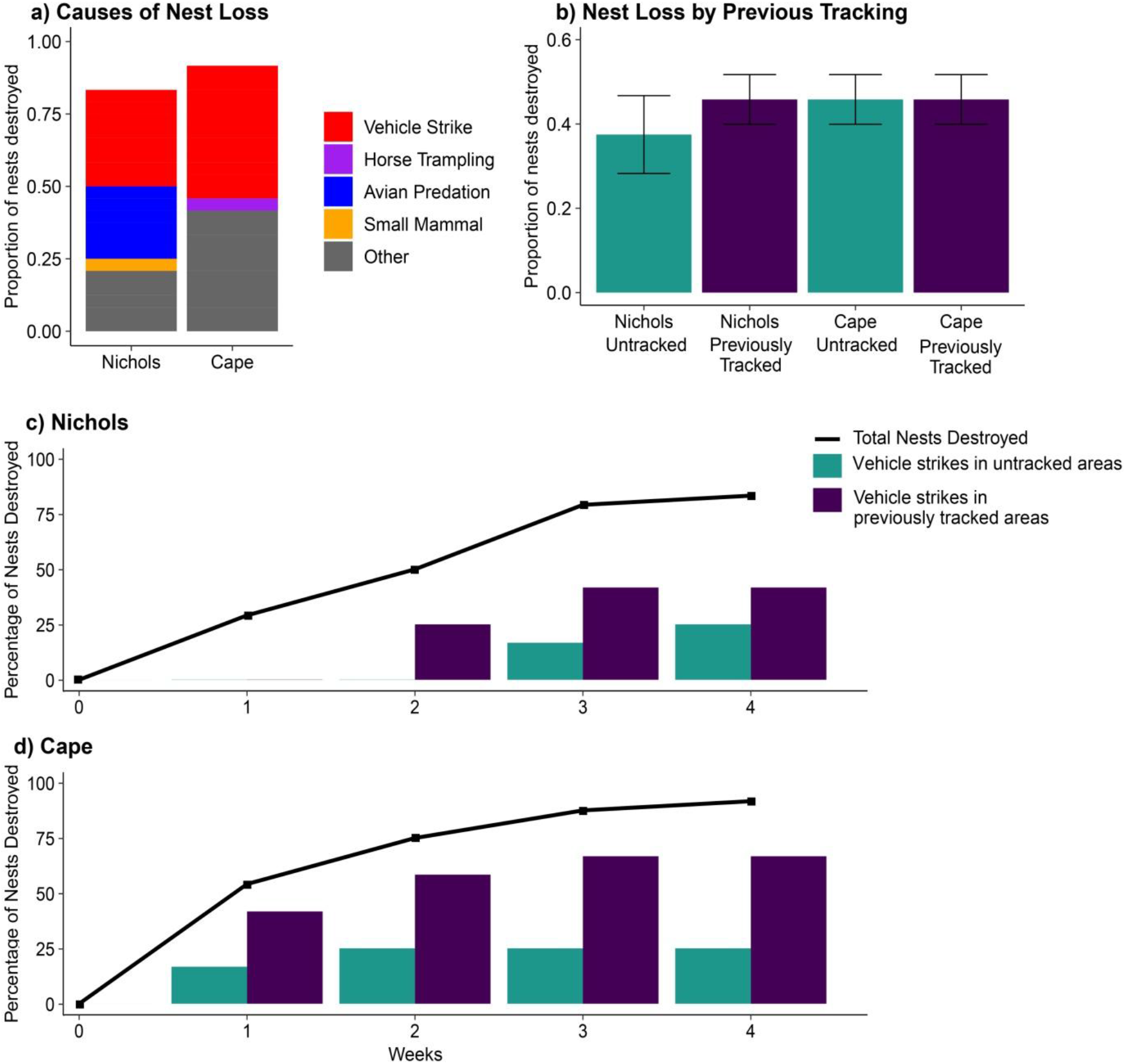
Artificial nest experiments at two study sites on the Marlborough coast where beaches have recently widened due to uplift from the 2016 Kaikōura earthquake. Four nests were located within each of six territories (n=24 nests per study site) and stratified by the presence of previous vehicle tracking (tracked / untracked) at the nest location. The Cape site is located near Cape Campbell where the high tide beaches are narrow. The Nichols site features wider high tide beaches and is located 4 km to the south (see Figure 1).

**Table 2.** Statistical comparisons for artificial nest experiments.

(a) Kruskal-Wallis tests on nest losses after 4 weeks.
| Independent variable | Dependent variable | $\chi^2$ | df | p† |
| --- | --- | --- | --- | --- |
| Territory | Total nest losses - all sites | 17.889 | 11 | 0.0841 |
| Territory | Total nest losses - Nichols | 5.75 | 5 | 0.3313 |
| Territory | Total nest losses - Cape | 10.455 | 5 | 0.06333 |
| Previous tracking status | Total nest losses - all sites | 0.8519 | 1 | 0.356 |
| Previous tracking status | Total nest losses - Nichols | 1.15 | 1 | 0.2835 |
| Previous tracking status | Total nest losses - Cape | 0 | 1 | 1.000 |
| Previous tracking status | Nest losses from vehicle strikes<br>- all sites | 4.1797 | 1 | **0.0409\*** |
| Previous tracking status | Nest losses from vehicle strikes<br>- Nichols | 0.7188 | 1 | 0.3966 |
| Previous tracking status | Nest losses from vehicle strikes<br>- Cape | 4.021 | 1 | **0.0449\*** |

|  | Estimate | SE | Z | p† |
| --- | --- | --- | --- | --- |
| Intercept | 1.1913 | 0.6039 | 1.973 | **0.0485\*** |
| Site | 1.4067 | 0.4855 | 2.897 | **0.0038\*\*** |
| Previous tracking status | 0.5538 | 0.5289 | 1.047 | 0.2950 |
| Site x Previous tracking status | 0.9806 | 0.7184 | 1.365 | 0.1722 |

|  | Estimate | SE | Z | p† |
| --- | --- | --- | --- | --- |
| Intercept | 1.2747 | 0.4273 | 2.983 | **0.0029\*\*** |
| Site | 0.9641 | 0.5901 | 1.634 | 0.1023 |
| Previous tracking status | 1.6272 | 0.4655 | 3.495 | **0.0005\*\*\*** |
| Site X Previous tracking status | 0.4317 | 0.7412 | 0.582 | 0.5602 |

† significant results at $\alpha = 0.05$ shown in **bold**. \* = $p < 0.05$ , \*\* = $p < 0.01$ , \*\*\* = $p < 0.001$

### 3.4 Dune responses

New dunes were observed along many stretches of coastline throughout the study area. Despite an abundance of new accommodation space, however, the speed and extent of new dune formation varied due to differences in the rate of vegetation establishment and interactions with sand supply. At the beach monitoring sites the width of the new dune zone varied from 10.5 m at Mussel Point to over 30 m at Aerial Beach and Long Point, and increased only slightly between the 2018 and 2019 summers (Table 3). The widest new dune zone (38 m) was recorded at Aerial Beach in 2019. This site has a relatively flat cross-shore profile and exemplified the general pattern of new dune development elsewhere involving sand deposition above the post-quake high tide line, often facilitated by vegetation establishment in the same areas. Two years after the earthquake, native sand-binders including spinifex (*Spinifex sericeus*) and knobby club rush (*Ficinia nodosa*) had established on many uplifted beaches along with invasive marram (*Ammophila arenaria*) and seasonal flushes of sea rocket (*Cakile edentula*).

**Table 3.** Dune monitoring results from cross-shore transects at three beaches uplifted by the 2016 Kaikōura earthquake.

| Study sites | Cross-shore width of new dune<sup>†</sup> (m) | New dune height<sup>‡</sup> (m) above MHWS<sup>†</sup> (mean ± SE) | Percentage vegetation cover<sup>†</sup> (mean ± SE) | Maximum vegetation height<sup>†</sup> (m) | Vehicle tracking density in new dune zone<sup>†</sup> (%) |
| --- | --- | --- | --- | --- | --- |
| *Mussel Point* |  |  |  |  |  |
| 2018 | 10.5 | 1.97 ± 0.19 | 24.08 ± 8.98 | 0.56 | 11.1 |
| 2019 | 13 | 2.13 ± 0.16 | 28.29 ± 7.87 | 0.65 | 15.5 |
| *Aerial Beach* |  |  |  |  |  |
| 2018 | 30.5 | 1.94 ± 0.16 | 9.00 ± 3.64 | 0.70 | 16.3 |
| 2019 | 38 | 1.95 ± 0.15 | 23.38 ± 4.73 | 0.75 | 30.3 |
| *Long Point* |  |  |  |  |  |
| 2018 | 31 | 2.15 ± 0.30 | 11.74 ± 3.87 | 0.54 | 32.6 |
| 2019 | 31 | 2.16 ± 0.29 | 12.79 ± 3.63 | 0.65 | 48.5 |

<sup>†</sup> average value from n=2 transects at each site
<sup>‡</sup> measured at 1 m increments on transect line, MHWS = Mean High Water Springs

Vegetation height and percentage cover increased at all monitoring sites but to a variable extent (Table 3). The largest increase of around 25% cover in new dune area was at Aerial Beach. The narrower Mussel Point site had the highest percentage cover (29%) associated with the establishment of a new belt of sand-binding vegetation above the old spring high tide line in front of the old dune toe (Figure 8A,B). In contrast, the Long Beach site had only sparse cover in the new dune zone (12-13%) that is characterised by a more sloping beach with mixed sand-gravel substrates, which provides a harsh environment for plant establishment. New dune formation at the site is partly dependent on vegetative growth from spinifex runners into the new space (Figure 8C-E). This vegetative growth is creating new sand and debris accumulations that will likely assist wind-blown recruits become established. However, the overall rate of vegetation colonisation appears to be slower than the on the finer-grain beaches at Aerial Beach and Mussel Point.

**Figure 8.**
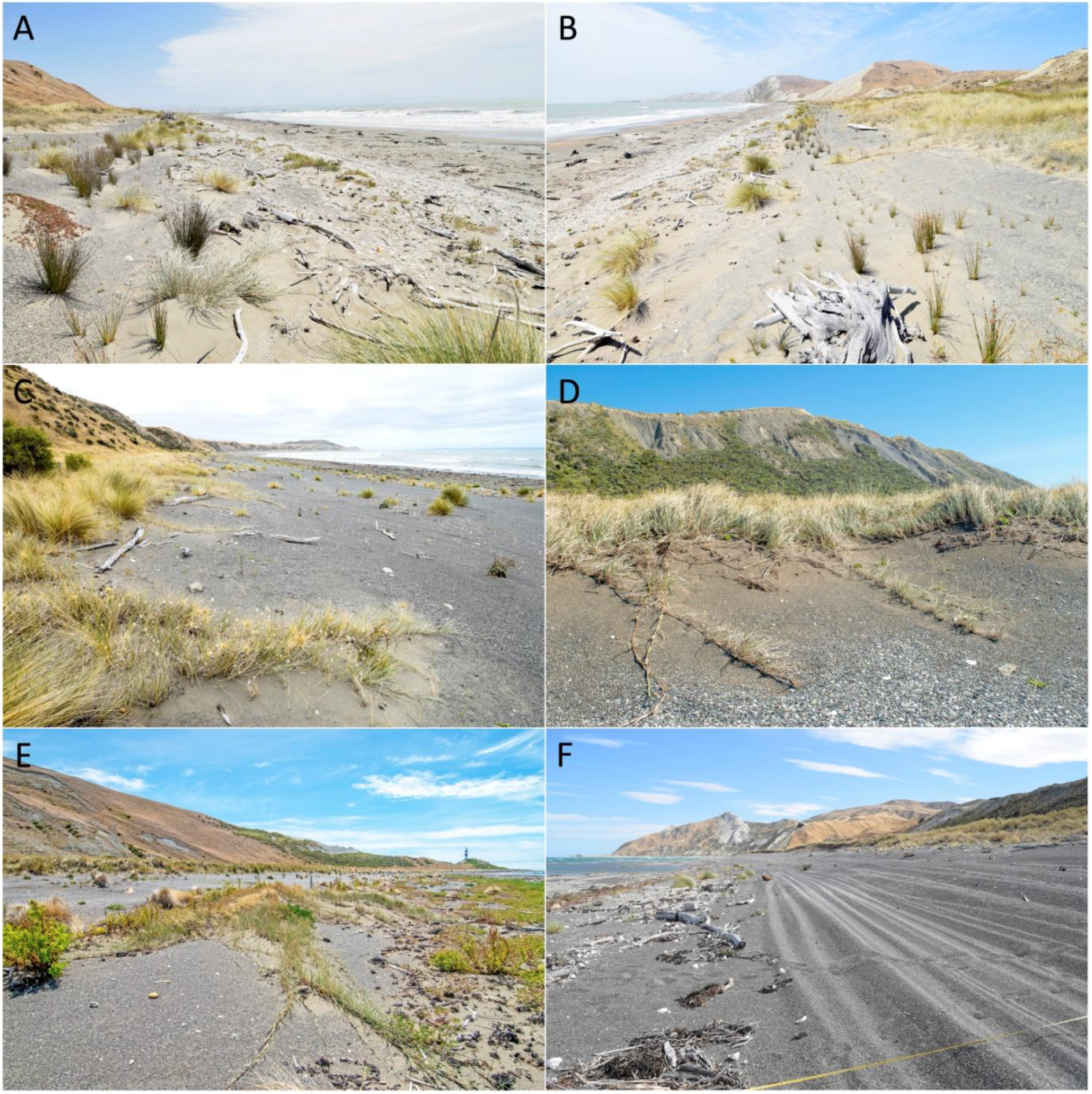
Dune monitoring sites. Mussel Point site with the new dune zone establishing seaward of old dunes looking west (A), and east (B). Spinifex runners from old-growth native dunes north of Long Point (C), and at the Long Point monitoring site (D). New plant establishment at Aerial Beach (E). Vehicle tracking in the new dune zone at Long Point (F).

Although some sand accumulation had already occurred in the new dune zone in the two years since the earthquake, further accumulation (as indicated by the average height of the profile), was only modest at all sites between 2018 and 2019 (Table 3). The observed vegetation cover and height increases did not appear to be sufficient to trap an appreciable quantity of additional sand between years. This may reflect nuances in the relationship between sand supply and the sediment-trapping properties of vegetation, or reflect erosion effects and other interactions.

The relatively slow rates of vegetation recovery and new dune formation since the earthquake are influenced considerably by people. Vehicle tracking densities, which provide a spatial measure of the extent of disturbance, were as high as 50% at Long Point by 2019 (Figure 8F), and increased at all sites between 2018 and 2019 (Table 3). The lowest tracking density (15%) at Mussel Point also coincided with the highest vegetation cover, suggesting a link between recent vehicle disturbance and the establishment of new plants. The tracking patterns were oriented mainly parallel to the shoreline, indicative of ORVs being used for long-shore access, but there were also examples of tracks on relatively steep dune faces with associated vegetation damage. At Mussel Point, for example, most of the ORV traffic utilises the intertidal beach area, and the increase in tracking in the new dune zone reflected a few vehicles (such as dune buggies) that were driven on sloping ground between the old dune toe and the new high tide line position. The tracking densities and trends measured at the monitoring sites were also similar to those observed elsewhere along the coastline (Figure 4). Greater increases in tracking density were observed at sites where there is a wider new dune zone that has become available for tracking. Reducing this footprint provides a focus for management to assist vegetation recovery and has potential to improve outcomes for native sand-binding species that are competing with more generalist exotics for the newly available area. Impacts on the growth of runners from existing native dune remnants were identified as a particular threat to be minimised where possible.

## 4. Discussion

### 4.1 Management of prograding shorelines

Tectonic uplift and isostatic adjustment from glacial de-loading are counterbalances to sea-level rise that will affect the speed and direction of coastal environment change (Church et al. 2013; Stammer et al. 2013). On the global stage, there are extensive coastal environments in seismically-active regions (Mogi 1974; Stern 2002), or subject to glacial isostasy (Peltier 2004; Shugar et al. 2014), and this suggests that a very considerable length of coastline is capable of progradation under conditions of uplift driven by these physical processes. Moreover, the contribution of uplift to shoreline evolution will remain important in the context of climate change since relative sea-level positions reflect the combination of vertical land mass motion and eustatic sea-level change (Cazenave & Llovel 2010). These factors contribute to variation in the magnitude and direction of relative sea-level changes at regional scales, with profound implications for day-to-day management (Nicholls & Cazenave 2010).

Erosion events and trends are commonly assessed as hazards with associated risks to assets and land-use (Crowell et al. 1999; Ferreira et al. 2006; Rabenold 2013), and the wealth of research effort associated with coastal erosion understandably reflects these concerns (Prasad & Kumar 2014). Although shoreline progradation effects such as beach widening are intuitively associated with benefits, our results show that they may also generate downsides. A key learning from this study is the need for rapid appraisals of new ‘opportunities’ and any potentially negative impacts of progradation events, much the same as is commonly exercised in response to erosion and shoreline retreat. Therefore, responding to the dynamics of shorelines in a timely fashion is the key principle that emerges.

Where patterns of resource use and values have been established in response to relatively stable conditions, physical landscape changes may present risks to the continuation of those values in place. In this case, deleterious effects were generated from the exercise of existing land-use rights, with the landscape having changed in relation to them. By contrast, there will often be beneficial opportunities associated with the creation of new land where previous erosion or the encroachment of infrastructure has exerted a coastal ‘squeeze’ (Doody 2004; Orchard et al. 2020a; Tono & Chmura 2013). For example, a reversal of coastal squeeze effects has been previously reported from uplift in the 2010 Maule earthquake in Chile (Rodil et al. 2015) and 2010-2011 Canterbury earthquake sequence in New Zealand (Orchard et al. 2020b). Additionally, sediment pulses originating from inland erosion and landslide events are known to exert pronounced effects on coastline evolution due to influences on sediment budgets at various spatio-temporal scales, and these can also lead to progradation events (Carter et al. 1987; Syvitski & Milliman 2007).

Anthropogenic responses to landscape changes, such as those reported here, must be cognisant of the requirements of natural ecosystems and their need to constantly readjust and reassemble in response to landscape change (Berry et al. 2013). The identification of their spatial requirements is therefore a key component of effective coastal conservation (Martinez et al. 2014). Coastal zonation patterns demonstrate that adjustments to relative sea-level positions are obligatory for many habitats and ecosystem types – yet they may take time to fully manifest following physical landscape change (Turner et al. 1998). These lag effects make the detection of change more difficult, but can be addressed using techniques such as spatial modelling to predict longer-term impacts, and longitudinal studies to confirm actual response trajectories and the outcomes that result (Orchard et al. 2020b; Watts et al. 2020).

In this case study, examples of natural ecosystem responses to shoreline progradation included the formation of new foredunes seaward of their pre-quake position. Although the associated spatial requirements were initially unknown, they became clearly evident one to two years after the earthquake as new vegetation became established on the uplifted shore. The winter storms of 2017 appeared to have had an equilibrating effect on the position of a persistent vegetation line, and in most places throughout the study area this line has changed little since. Additionally, there has been increasing vegetation cover landward of this line which now forms the new dune toe (Table 3). Future changes not directly observable within the time frame of this study may include the landward boundary of the dune system becoming more stable as the new foredune intercepts sand supply. The likely consequence is progressive invasion of the sand-binding species by other terrestrial vegetation types. Consequently, the beach widening changes associated with tectonic uplift are expected to cause a seaward shift in the entire dune system over time. This illustrates principle of habitat migration in which, despite the nuances of lag effects, entire systems must move to new locations if they are to persist under conditions of change (Hovick et al. 2016; Schlacher et al. 2008b).

In the case of banded dotterel nesting habitat, there was an expansion of the sparsely vegetated substrates that are required for nesting success (Pierce 1983; Rebergen et al. 1998). However, plant community succession is also well underway on the uplifted gravel beaches (that support the majority of nesting habitat), in addition to the dune responses discussed above. This illustrates an interaction between vegetation recovery and nesting habitat, and in the absence of other changes is expected to progressively reduce the area of suitable habitat. In the meantime, the apparent bonus effect of additional habitat has been counterbalanced by an increase in ORV use that poses a considerable threat to nesting (Figure 7). Similar ORV impacts have been reported in a study of hooded plover (*Charadrius rubricollis*) with a vehicle tracking density of 20% that resulted in an average of 6% of artificial nests being run over per day, implying an 81% loss within the incubation period (Buick & Paton 1989). In our case, these new human pressures are similarly un-managed at present, but were facilitated by physical landscape changes that rapidly altered the pattern of recreational activities rather than being the result of gradual access and user group changes that are typical of other studies (Luke & Schlacher 2008; Priskin 2003). In all likelihood, the new threats from ORV traffic have negated any potential reprieve from predation pressures conferred by the widening of high tide beaches, and demonstrate an important anthropogenic dimension for planning following natural disaster events.

### 4.2 Integrating recreation and conservation

In theory, there is potential for win-win outcomes from the landscape changes reported here, if low impact options for ORV access could be identified using techniques such as designated routes. In practice, however, this may be difficult to achieve due to the widespread distribution of fragile vegetation types and shorebird nesting grounds that in many ways are a reflection of the high biodiversity values of this formerly-remote coast. This presents a conundrum for the design of fine-scale spatial planning approaches that might be used to weave a vehicle route through the new landscape now that the major physical barriers, such as headlands, are more easily bypassed. Three important dimensions for the identification of an integrated solution are discussed in the sections below, before summarising key learnings from this case in relation to the policy context.

#### Designated vehicle routes in intertidal areas

Due to the distances that may be travelled in daily ORV excursions in our study area, a reliance on intertidal (e.g., low-tide) vehicle travel routes may not be an effective management solution for the protection of high tide beaches, since vehicle movements are likely to occur in those areas on return journeys. Additionally, there may also be impacts associated with ORV use in intertidal areas, since crushing effects have been reported for a wide range of sandy beach infauna (Lucrezi & Schlacher 2010; Moss & McPhee 2006; Schlacher et al. 2008a; Schlacher et al. 2008c). New Zealand examples include impacts on shellfish such as toheroa (*Paphies ventricosa*) and tuatua (*Paphies subtriangulata*), and include sensitive intertidal elevation zones related to the distribution of different size-classes (Moller et al. 2014; Taylor et al. 2012; Williams et al. 2013). Consequently, the mapping of sensitive intertidal areas would be required to establish the degree of impact associated with ORV use and any potential low-impact options (Schlacher & Thompson 2007).

#### Designated vehicle routes at the top of the beach

Conceivably, a low-impact vehicle route could be found at the top of the beach if it represented a less sensitive environment. However, the degradation of old dune systems, that are characteristic of this location, is not a permissible activity due to their conservation status as endangered ecosystems (Holdaway et al. 2012). Aside from their existing conservation value they are crucial for progression of the current recovery process as the seed sources for new recruitment. Additionally, in the case of the primary sand-binding species *Spinifex sericeus,* repair of the dune system is facilitated by the vegetative growth of runners from the dune toe (Bergin 2008; Orchard 2014), and vehicle tracking in these areas are a further threat to recovery and seaward dune migration. Overall, the concept of a formed roadway in stable and less sensitive areas that may be found inland of coastal dunes may be appropriate when and where those conditions exist, but such a route would lie outside of the current dune system.

#### Designated vehicle access areas for key activities such as boat launching

An alternative set of options for the consideration of ORV access with appropriate environmental protection involves the identification of designated vehicle *areas* (being sections of coast where ORV use is permitted) rather than routes through sensitive areas. Such areas might also include a few ‘sacrificial’ activity zones where high impact activities such as driving on dunes is permitted as a strategy to reduce impacts elsewhere; and these would logically be situated in areas where such impacts are already occurring. This line of thinking also applies to the continued use of vehicles at key sites for activities such as boat launching, especially where these activities are already occurring (and indeed may have a long history in some places). In relation to the previous management context, it is important to recognise that the post-disaster setting presents a new set of considerations that require active management to enable the continuation of such uses in an appropriate manner. These management responsibilities arise because of the increased potential for problematic ORV access into adjacent coastal areas associated with the landscape changes, leading to the need for formal controls or effective non-statutory interventions to ensure that such problems do not occur. Practically speaking, this means that coastal managers must designate vehicle access areas and their boundaries, in contrast to the pre-quake situation in which there were no formal provisions for appropriate locations and their use. While the contemplation of designated vehicle routes through sensitive areas depends heavily on the results of impact assessment work, the identification of vehicle access *areas* at key locations provides a tangible and important starting point.

### 4.3 Responses and responsibilities under the policy context

Impact assessments are a required aspect of the environmental policy cycle in New Zealand and similarly in other jurisdictions. For example, knowledge of impacts is typically necessary for day-to-day decisions on audits or permits functions, and in the review of statutory policies and plans at local through to national levels. A hierarchical arrangement of these measures is required by the Resource Management Act 1991 (RMA) which is New Zealand’s primary environmental legislation alongside the Conservation Act 1987, Wildlife Act 1953, and Reserves Act 1977 (and amendments) (Memon & Perkins 2000). National Policy Statements prepared under the RMA are a key tool for environmental management that direct the preparation of more specific policies and plans within a devolved organisational framework (Memon 2002). Under the New Zealand Coastal Policy Statement 2010 (NZCPS), requirements for the protection of coastal environments include avoiding adverse effects on indigenous species, ecosystems and vegetation types that are listed as threatened or at risk, or are naturally rare, and others that are at the limit of their natural range, or contain nationally significant examples of indigenous community types. Significant adverse effects must also be avoided in indigenous ecosystems and habitats that are only found in the coastal environment and are particularly vulnerable to modification, including estuaries, lagoons, coastal wetlands, dunelands, intertidal zones, rocky reef systems, eelgrass and saltmarsh; and similarly for areas of predominantly indigenous vegetation, and habitats that are important during vulnerable life stages, or for recreational, commercial, traditional or cultural purposes (Department of Conservation 2010). In practice, implementation of this statutory framework creates a substantial obligation for environmental planning informed by baseline and impact assessments (Orchard 2011). Management authorities, such as regional and district councils, are legally required to fulfil the statutory responsibilities set out in higher level legislation, creating a form of decentralised governance that is also common in other countries (UNDP 2004).

In addressing NZCPS requirements (and other legislation), methods to avoid the adverse effects of ORV use present a considerable challenge for management authorities. In remote settings where the enforcement is difficult, non-statutory measures that incentivise desirable driver behaviour are likely to be beneficial, alongside regulatory tools. For example, the support of user groups for spatial planning measures, such as permitted areas or vehicle routes, could play a self-reinforcing role by helping to socialise and generate buy-in for effective solutions. Conversely, there is an opportunity for user groups to develop initiatives around the design of such systems in advance of regulatory measures. Previous studies have shown that the success of motivational incentives also depends considerably on relationship-building between the public service, non-governmental organisations and local community stakeholders (Gunningham 2009; Koontz et al. 2004).

The establishment of areas that are closed to vehicles is one of the most obvious options for preventing their impacts on sensitive species and habitats. Moreover, beach closures at the location of shorebird nesting grounds have been shown to be effective in studies from Australia (Weston et al. 2012) and Namibia (Braby et al. 2001). In addition to reducing direct effects on nesting success, such as egg-crushing, protected areas can assist with reducing disturbance effects on essential wildlife behaviours such as incubation and foraging (Ruhlen et al. 2003; Weston & Elgar 2007). In combination, these considerations can inform decisions on the size and configuration of protected areas to optimise their efficacy and ensure they are well integrated with public access opportunities. These strategies, behaviours, and good practice examples can be incentivised by non-statutory motivational initiatives led by either NGOs or government authorities, and are a great focus for collaborative design and promotion. If such measures prove to be effective at reducing impacts where vehicle access is permitted, they will undoubtedly improve its consistency with established environmental objectives.

### 4.4 How can beach users reduce their impacts?

While management authorities must respond to their statutory obligations under relevant legislation, there are certainly voluntary actions available to beach users to reduce their actual and potential impacts. They include, perhaps most importantly, the use of existing vehicle tracks where they exist, and the exercise of caution or reconsideration if there are no such well-formed tracks. When travelling in environmentally sensitive areas, ORV users can take practical steps such as looking out for wildlife, maintaining an appropriate separation distance, taking note of fragile vegetation and habitat types, and keeping an eye out for signage and various types of exclosures that can be used to indicate their presence. Driving speeds and setback distances from wildlife are known to be important determinants of the degree of disturbance associated with motorised vehicles (Schlacher et al. 2013). By extension, other characteristics of vehicle types, such as the level of noise they emit, are also likely to contribute to the adverse impacts of vehicle use in particular situations. Conversely, this presents opportunities for vehicle users to select their transport mode purposefully to avoid impacts and gain the acceptance of other stakeholders. In the near future, for example, electric bikes and other electric vehicle types may facilitate less invasive access modes in sensitive wildlife habitats, improving the options for sustainable travel in these areas. Ultimately a consistent focus on recreational impacts could be applied to all access modes to help manifest a light-footed, low impact relationship with the natural environment while encouraging visitation by people.

### 4.4 Concluding remarks

This study highlights the need to provide for natural environments in responses to landscape changes. Failure to do so is likely to undermine conservation objectives with potentially drastic consequences. Even small changes in critical parameters can lead to new impacts over large areas. Recovery actions following disturbances must pay attention to impacts from both the original disturbance and consequential changes, and identify trade-offs between desirable objectives. This study illustrates these requirements in a situation of shoreline progradation that offered new land-use opportunities for various forms of nature-based recreation and access. The recreation-conservation nexus was a key driver of change and negative impacts were generated by the interaction between access opportunities and dynamic natural environments. A key lesson that emerges is the need for timely impact assessments across the social-ecological spectrum whenever physical landscape changes alter the accessibility of geographical locations and resources. Similar principles are transferable to many other natural hazard and disaster recovery settings and will help to promote the achievement of sustainable development and resource management goals.

## 5. Acknowledgements

This research was funded by the Ministry of Business Innovation and Employment (MBIE) Endeavour Fund (UOCX1704), with additional support from Ministry for Primary Industries (MPI) (KAI2016-05) and Sustainable Seas National Science Challenge (UOA20203). We thank our research partner, Marlborough District Council, for supporting this project, and the East Coast Protection Group for their continuing interest. Thanks to Thomas Falconer, Zoe Smeele, Ben Crichton, Ryan Taylor, Eva Pomeroy, Shelagh Taylor, and Keith Jacob for field assistance, and John Pirker for iwi liaison. Particular thanks to local landowners, especially Sally, Rob and Thomas Peters, Kevin Loe and community members who have assisted with survey logistics and the establishment of study sites in support of this project. The research reported here also benefited from helpful conversations with a wide range of governmental and non-governmental groups and interested individuals.

