## Supplementary Material for "Managing beach access and vehicle impacts following reconfiguration of the landscape by a natural hazard event"

---

**Figure S1** Examples of motivational signage to protect sensitive ecological areas from human disturbance on beaches. (A) and (B) Australia, (C) and (D) New Zealand.

**Figure S2** (A) Typical example of a banded dotterel (*Charadrius bicinctus bicinctus*) nest on the high tide beach. The location is Canterbury Gully. (B) Banded dotterel chick. (C) and (D) Examples of artificial nest experiments at Nichols and Cape Campbell. (E) Peck marks characteristic of avian predation. (F) Nest crushing from a vehicle strike in the artificial nest experiment.

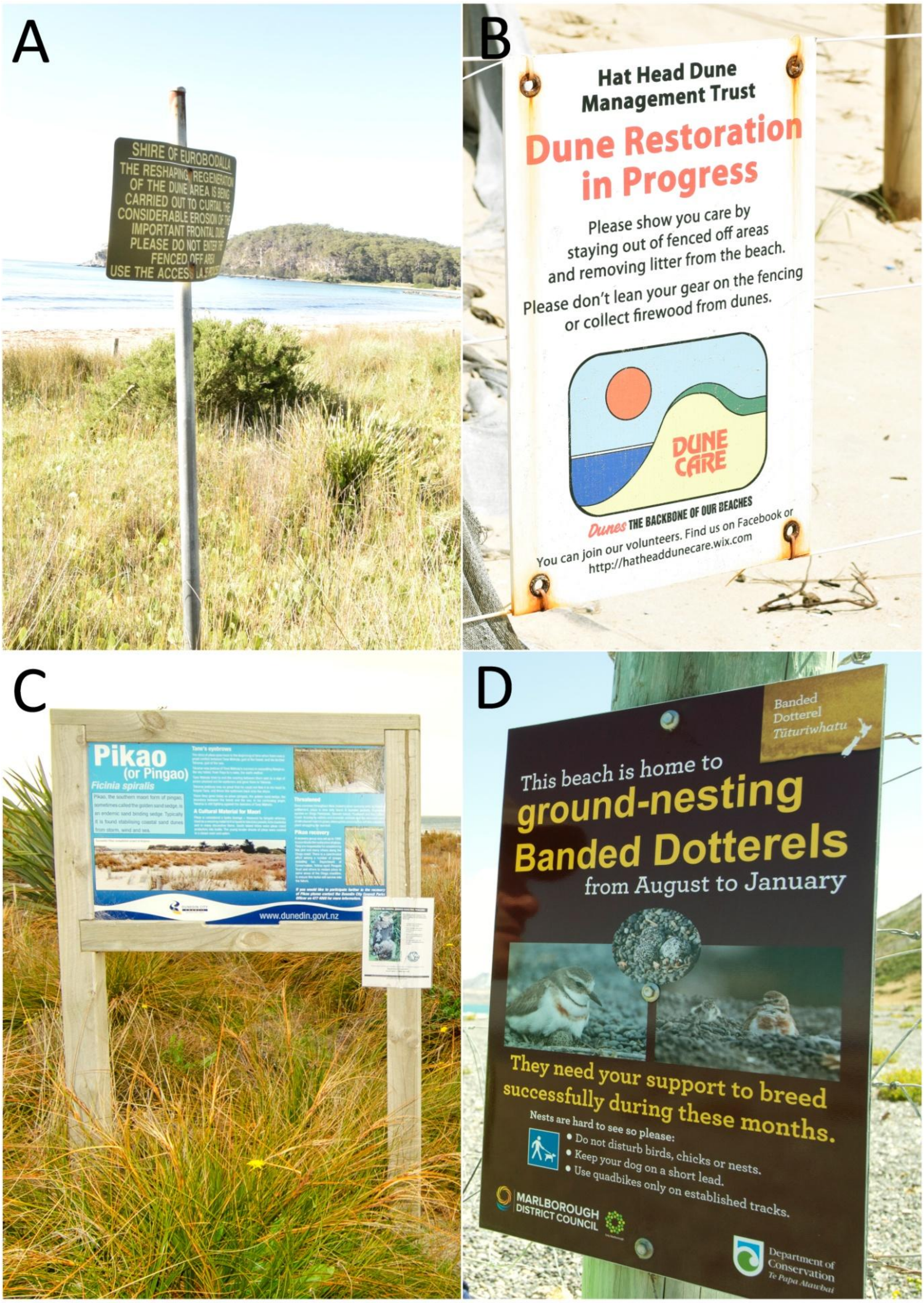


**Figure S1** Examples of motivational signage to protect sensitive ecological areas from human disturbance on beaches. (A) and (B) Australia, (C) and (D) New Zealand.

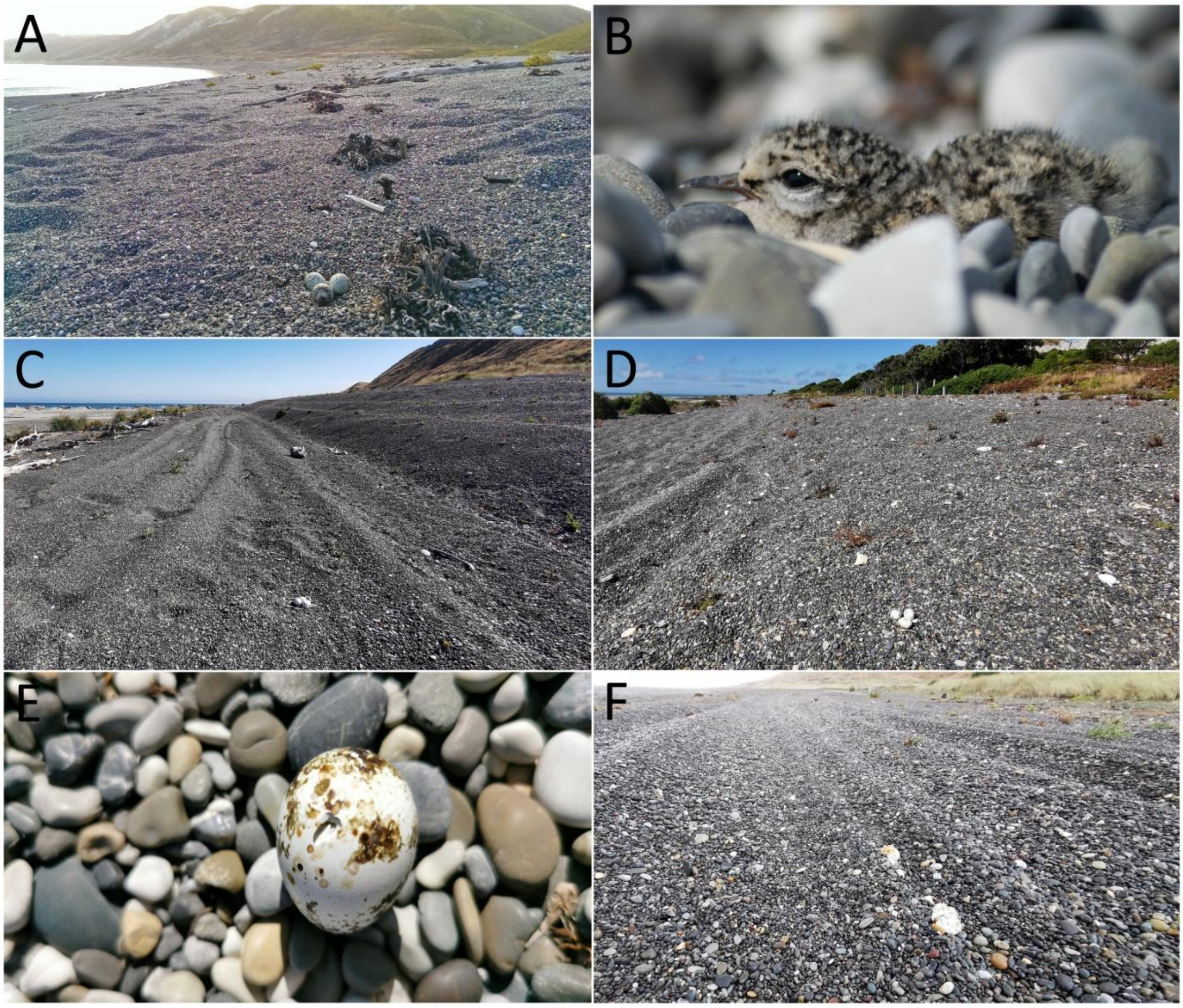


**Figure S2** (A) Typical example of a banded dotterel (*Charadrius bicinctus bicinctus*) nest on the high tide beach. The location is Canterbury Gully. (B) Banded dotterel chick. (C) and (D) Examples of artificial nest experiments at Nichols and Cape Campbell. (E) Peck marks characteristic of avian predation. (F) Nest crushing from a vehicle strike in the artificial nest experiment.
